# Grasping Under Target Size and Location Uncertainty

**DOI:** 10.64898/2026.10.08.757378

**Authors:** Kevin A. LeBlanc, Anne T.D. Lacroix, Jada A. Benwell, McKenna Grant, Brett B.T. Feltmate, Heather F. Neyedli

## Abstract

Everyday actions often require movements to be planned before the exact target is known, yet most research on grasping has examined situations in which individuals reach toward a single, well-defined object. The present study investigated how grasping actions are planned under target uncertainty using a go-before-you-know (GBYK) paradigm, in which participants initiated their movement before knowing which of two potential objects would become the target. Using 3D motion capture, we examined whether peak grip aperture (PGA) was influenced by the size of a competing nontarget object. Participants grasped cylindrical objects that varied in size under two conditions: GBYK, in which target identity was revealed after movement initiation, and know-before-you-go (KBYG), in which the target was known prior to movement onset. Across two experiments, participants appropriately scaled grip aperture according to the target object in both conditions. However, during GBYK trials, PGA was influenced by the presence of a larger nontarget object. Specifically, when grasping a small target paired with a large nontarget, participants produced larger grip apertures, whereas a small nontarget did not influence grasping toward a large target. This asymmetric interference pattern is inconsistent with a motor averaging account, which predicts bidirectional shifts toward an intermediate aperture. Instead, the findings suggest that under conditions of uncertainty, grasp planning incorporates competing object information in a manner that prioritizes successful object acquisition. Together, these findings extend research on action planning under uncertainty beyond reaching and highlight the dynamic nature of grasp planning when multiple potential targets are present.

**NEW & NOTEWORTHY:** This study provides the first examination of grasp planning under target uncertainty using a go-before-you-know paradigm. Unlike previous work focused primarily on reaching movements, we show that grasp formation dynamically incorporates information from competing potential targets. Under uncertainty, grip aperture was biased by larger nontarget objects in a manner consistent with prioritizing successful object acquisition rather than adopting an intermediate grip aperture between competing motor plans.

## INTRODUCTION

While there has been substantial scientific progress towards understanding the neuroscience of grasping (1–6). Much of the previous research has focused on paradigms where participants are simply required to reach out and perform a grasping action to one single target object. This presents a limited view of grasping because we accomplish most of our daily tasks by performing actions in the presence of multiple possible target objects. Yet it remains unclear on how the brain plans the kinematics of a grasping action when multiple potential targets are present. Furthermore, interaction in the real-world involves continuous updating of motor plans in response to incoming sensory information. Often actions need to be planned rapidly under uncertainty about the true state of the world. This contrasts to the linear stimulus, react, response nature of typical trials in experimental psychology.

Within the past decade, a paradigm termed ‘go-before-you-know’ (GBYK) has emerged in which participants must initiate their movement before the target location is revealed while multiple potential target locations are present, allowing researchers to investigate how actions are planned under conditions of uncertainty. Findings show that participants often initiate a reaching movement that is intermediate between potential target locations (7–8). Initially, it was proposed that the intermediate movement was a result of the partially activated action representations associated with the potential targets being averaged together (9–12). However, subsequent evidence favours a more dynamic view in which a single movement is planned to optimize task success, and the plan is then updated once target information becomes available (13–17).

The vast majority of the GBYK studies have focused on reaching and pointing movements where the primary outcome reflects the transport of the limb toward a target (7–13, 16–17). Although the GBYK paradigm has also been tested with saccadic eye movements (19) and stepping actions (18), the GBYK paradigm has yet to be applied to grasping, an action that extends beyond limb transport by requiring coordinated control of grip formation and object manipulation. Specifically, grasping can be divided into three components: transporting the hand toward the object, shaping the fingers to match the object’s properties (e.g., size and orientation), and manipulating the object once contact is made (20). During reach-to-grasp movements, the hand is initially transported rapidly toward the target object, followed by a slower terminal phase in which the fingers close around the object (21–22). Grip formation begins early within the movement, with fingers progressively opening to a peak grip aperture (PGA). This identifiable landmark of the movement is highly correlated with the size of the target object (20, 23–24). Unlike reaching movements, grasping permits examination of both the transport component of the action and the evolution of grip formation, providing a unique opportunity to determine whether action uncertainty extends beyond where the hand is moving to include how the hand is shaped for object interaction.

Since the hand starts to reflect the size of the target object as soon as a reach-to-grasp movement is initiated, this reflects that the visuomotor system incorporates object information from the earliest stages of action planning. At the neural level, converging evidence from monkey neurophysiological and human neuroimaging studies have shown that grasping relies on a distributed parieto-frontal network that transforms visual object information into motor commands required to successfully perform a grasping action (3, 24–31). Visual information relevant for performing a grasping action such as, size, shape, orientation, and weight are represented across both dorsal and ventral visual streams (32–41). However, the dorsal stream is primarily responsible for transforming visual information into the visuomotor representations necessary for planning visually guided grasping actions (29, 42).

These distributed representations are integrated through interactions between parietal and frontal regions to generate appropriate motor commands for object-directed actions. The anterior intraparietal sulcus (AIP) plays an important role in this network by integrating visual information about object properties and transmitting this information to premotor regions responsible for specifying grasp configurations (43–45). Neurophysiological evidence further suggests that the ventral premotor cortex (PMv) primarily contributes to hand shaping, whereas the dorsal premotor cortex (PMd) is more closely associated with grip and wrist orientation (46–49). Collectively, this parieto-frontal network enables grasping movements to be rapidly adjusted based on the physical properties of objects, producing remarkably accurate and efficient actions under typical laboratory conditions. However, much of this evidence comes from paradigms in which participants perform a single action toward a single, well-defined target. This differs substantially from everyday behaviour, where multiple potential targets and competing action possibilities are often present. Therefore, an important question is how the grasping network resolves competition between potential actions and configures grasp kinematics when multiple action options are available.

In non-human primates, it has been shown that prior to knowing which of two grip types to use for a single object, the motor cortex encodes multiple action representations in parallel (50), similar to the parallel encoding for reach direction observed for reaching movements in the premotor cortex (51–52), and these representations can then combine into a single optimized plan that accounts for both potential grips (15). However, this evidence is limited to a single object affording both grip options. Thus, it remains unknown how the brain plans the kinematics of a grasping action when multiple potential targets are present, and movement is initiated prior to target selection resulting in target uncertainty.

It is important to note that previous studies have shown that the shaping of hand aperture during a single grasping action, when target is known and under on-line control, is typically very resistant to interference from distractor objects (53–54). However, when attention is directed toward distractor objects (55–56), or when grasping is performed in the presence of multisensory distractors (1), evidence suggests that distractors can influence grasp kinematics. Building on this, the present study takes an exploratory approach to investigate how (or whether) grip aperture can be influenced by the size of nontarget objects when planned under uncertainty within the context of a GBYK paradigm.

Using 3D motion capture to measure hand kinematics, participants were presented with two cylindrical objects (small or large), and required to grasp one of them under two different conditions. For the GBYK condition they initiated their grasping movement before knowing which object was the target, and for the KBYG (know-before-you-go) condition the target object was indicated prior to movement onset. Within both conditions, objects on each trial were either the same size or differ in size. Of particular interest are the trials where the objects differed in size to explore the impact the nontarget object has on PGA. In addition, some participants started with their index finger and thumb together (experiment 1), while others started with them apart (experiment 2), allowing us to more precisely examine how grip scaling is influenced by the size of the nontarget. Based on the “digit-in-space” theory of grip formation (57), simultaneous presentation of differently sized objects could create overlap in the aperture tuning required for each object, and the starting finger position may modulate this effect.

Four possible patterns of results may arise in the GBYK condition, compared to the KBYG condition, which is expected to produce the target-appropriate grip scaling unaffected by the nontarget:

- Symmetric intermediate movement hypothesis: An intermediate grip aperture is produced when the two objects differ in size, resulting in a smaller grip aperture when the large object is the target with a small distractor, and larger grip aperture when the small object is the target with a large distractor, resulting in a symmetric pattern of grip aperture modulation. This pattern would be consistent with motor averaging, whereby the competing object representation is integrated to produce an intermediate grip aperture. Such averaging could reflect a form of strategic optimization, in which participants adopt an intermediate grip aperture to maximize the likelihood of successfully grasping either potential target.
- Safety optimization hypothesis: Grip aperture defaults to the largest object to ensure a successful grasp (e.g., reaching with a larger aperture for the small object still ensures a successful grasp, whereas reaching with smaller aperture for the larger object can risk tipping the object over), such that the presence of a small distractor would not influence grip aperture when reaching for the large object, but the grip aperture would be larger when the small object is the target, and the large distractor is present. This pattern would reflect a safety-based strategy in which participants prioritize avoiding grasp failure associated with underestimating object size, even if this results in a larger-than-necessary grip aperture when grasping the smaller object.
- Precision-first hypothesis: Grip aperture defaults to the smallest object since it demands more precise on-line visuomotor control (58). In this case, the presence of a small distractor would cause a smaller aperture when the large object is the target. However, the presence of large distractor would not affect the aperture towards the small object.
- Overlapping mechanisms hypothesis: Grip aperture may be influenced by the starting position of the fingers, reflecting overlapping constraints in aperture scaling. Specifically, when participants start with their fingers together, the aperture tuning for either object must pass through the aperture required for the small object, producing interference from a small distractor but not a large one (see 59 for evidence of this pattern in a sequential grasping task). Conversely, when starting with fingers apart, the aperture tuning must pass through the aperture required for the large object, leading to interference from a large distractor but not a small one.

These experiments will provide a novel test of how the nervous system plans grasping actions under uncertainty and offers a new way to examine how competing action possibilities are resolved during the early stages of motor planning.

## MATERIALS AND METHODS

### Experiment 1

#### Participants

An a priori power analysis indicated that a sample size of 23 participants per experiment was required to detect a medium effect size (partial η² = .06). Accordingly, twenty-three students (17 female, *Mage* = 19.4 years, ± 2.) at Dalhousie University participated in the current study in exchange for partial course credit. All participants had normal or corrected-to-normal vision and no history of neurological deficit as ascertained by self-report. Because all grasping movements were performed and measured using the right hand, participants were required to be comfortable performing grasping actions with their right hand. Nineteen participants reported being right-hand dominant, while four reported being left-hand dominant. Each participant provided informed consent prior to participation in accordance with guidelines established by the Dalhousie University Research Ethics Board.

#### Materials

An OptiTrack 3D motion capture system using eight PrimeX 13 cameras (120Hz sampling frequency) was used to collect data for live kinematic capture of the participant’s right hand. OptiTrack PrimeX 13 cameras captured light from ten infrared markers - eight active (OptiTrack Active Tag) and two passive - to record the coordinates of the marker’s x, y, and z locations. The markers were positioned on the participant’s right hand (Figure 1). A python-based framework (KLibs) was used to run the experiment program, using the software Motive for live input of the marker coordinates during movement. Data preprocessing, statistical analysis, and figure making was completed in R.

**Figure 1.**
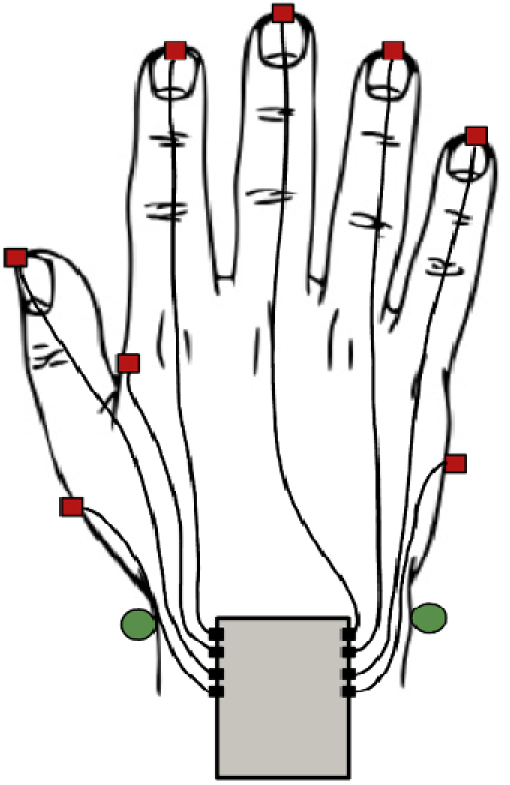
Infrared marker placement on participant’s right hand: Passive markers indicated as circles (green). Active markers are indicated by squares (red). Markers and their respective components are secured to the participant to prevent interference with the OptiTrack cameras. Markers were placed as follows: medial-ulnar aspect of the wrist joint (passive), lateral-radial aspect of the wrist joint (passive), one marker situated at the distal end of each phalange (active), proximal and lateral to the first metacarpophalangeal joint (active), proximal-medial to the first metacarpal and proximal-lateral to the second metacarpal (active), medial boarder of the hand half way between the fifth metacarpophalangeal joint and the wrist joint (active).

A 27” ViewSonic monitor was used to display target selection stimuli. 3D printed white cylinders with a height of 5 cm and a diameter of either 4 cm (small) or 8 cm (large), with a weight of 33.6 g or 128.6 g respectively, were used as potential targets. Each cylinder was padded with sound dampening felt on the bottom to prevent the participants from gathering audio feedback during experiment set up.

Liquid-crystal display (LCD) occlusion glasses (PLATO Translucent Technologies, Toronto, ON, Canada) were worn by the participant for the duration of the experiment to allow for control of stimuli presentation. In their resting state, LCD glasses are opaque, blocking the participant’s view. When activated by the experiment program at the start of each trial, the glasses turned transparent, allowing the participant to fully see the targets and perform the grasping task.

#### Procedure

Prior to beginning the experiment, the table surface was set to the participant’s waist height, and participants entered their demographic information into the program and were given a task explanation prior to initiating testing.

For each trial, on the table surface was a keyboard facing the participant and a centered monitor lying flat on the table in front of the participant. The starting location was the space bar marked by tape to provide tactile feedback. Two cylinders were placed on top of the screen 30 cm in front of the starting location (center of the space key), at a 30° angle from the midline. The target object was illuminated by presenting a white circle 1 cm larger than the object’s diameter on the screen underneath it (Figure 2).

**Figure 2.**
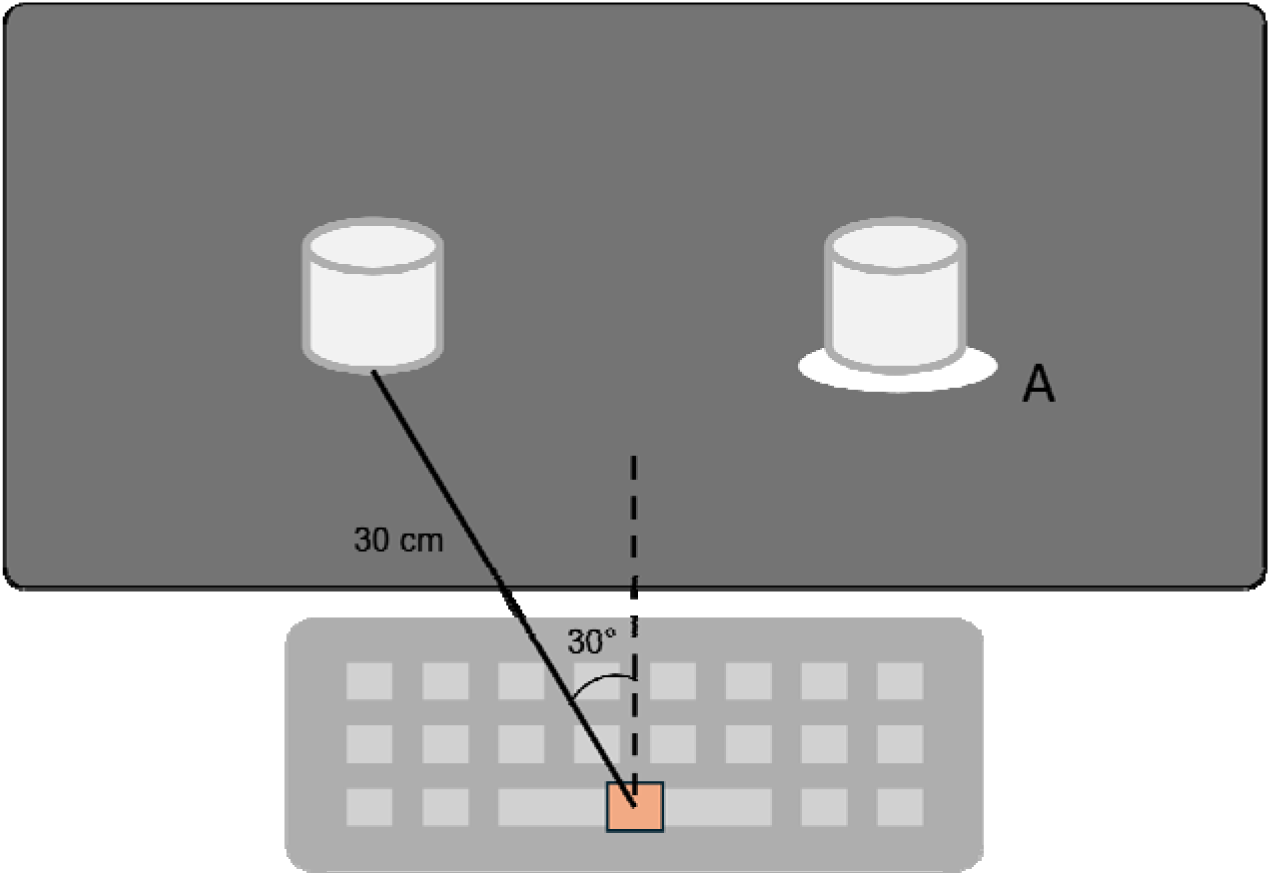
Stimulus layout. Start location is at the center of the space key on the keyboard (represented by the orange box). Each object is 30 cm from the start location and at a 30-degree angle from the midline. Target was indicated by illuminated a circular outline underneath the object (A). In this example, the target would be the object on the right.

There were two conditions, each consisting of eight practice trials followed by 80 test trials (160 total test trials). Because the GBYK condition required participants to initiate movements under target uncertainty rather than toward a known target, participants always completed the GBYK condition first to familiarize themselves with this novel task demand, followed by the KBYG condition, with an optional break of 1-3 minutes between conditions. Target object size, distractor object size, and target object location were randomized for each trial using KLibs wherein 10 trials per combination were presented.

The LCD glasses were opaque at the start of each trial while the experimenter positioned the objects. Once the objects were correctly positioned, the participant would trigger the start of the trial by holding down the space key at the starting location with right index finger and thumb pressed together.

At the start of the trial the LCD glasses turn transparent, followed by the presentation of 784Hz sine-wave tone lasting 100ms, after a randomized delay time of 500-2000ms to prevent expectancy effects, signaling movement onset. In the GBYK condition, the target was only illuminated once the participants moved their hand a randomized distance between 5-10cm from the starting position to prevent expectancy effects. In the KBYG condition, the target was illuminated prior to the LCD glasses turning transparent, before the participants began movement. Participants were required to lift the target object from above using their thumb and index finger as a precision grip (Figure 3). Participants had 1000ms after the audio tone to complete their grasping movement prior to the LCD glasses returning to resting state.

**Figure 3.**
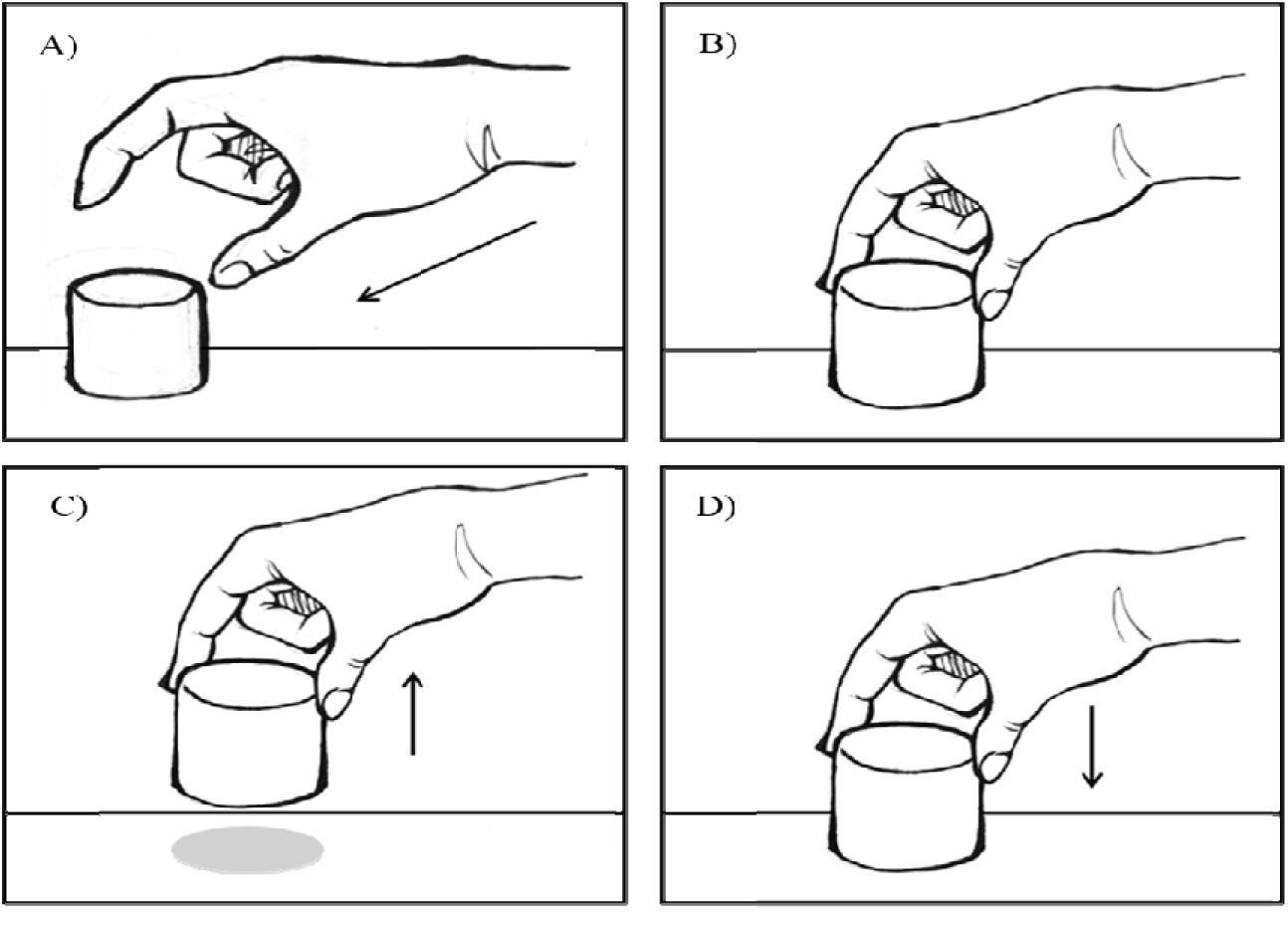
Visual depiction of participant’s instructed grasping movement: (A) Participant reached for the target object. (B) Participant grasped the target object from above, using only the thumb and index finger of their right hand. (C) Participant lifted the object. (D) Participant placed the object back down in the same location.

A trial that exceeded the time limit or contained a false start (early release of the space key) was considered null and was recycled, not counting towards the 160 ‘successful’ test trials.

#### Data Processing

The raw 3D-position data of the infrared markers were processed online and filtered using a fourth order dual low-pass Butterworth filter with a cut-off frequency of 6Hz. Grip aperture was measured in millimeters as the Euclidean distance between the markers of the thumb and index finger. Hand position was measured as the mean x, y, and z positions of the hand markers (excluding finger markers), wherein the starting hand position was indicated as the origin. Movement start was defined as when the participant released the space key, and movement end was defined when the participant began lifting the object.

Trials were excluded if the starting grip aperture or initial hand position was ± 3 SDs from individual participant’s mean values (5.7% of trials removed). As trial duration varied, time was normalized using a spline transformation ensuring every trial consisted of 1000 frames, providing a time standardized dataset.

A linear mixed-effects model was conducted to examine the effects of target size, distractor size, and target location on peak grip aperture separately for the GBYK trials and the KBYG trials. Alpha was set at 0.05 for main effects and interactions. Because the impact of distractor size was part of our hypotheses, we a-priori planned to compare PGA between the small and large distractor trials using Bonferroni corrected LMEs for each combination of conditions within the LME (e.g., each combination of target location and target size). Between the GBYK and KBYG trials this yielded a total of 8 potential comparisons between distractors sizes for PGA resulting in a Bonferroni corrected alpha of 0.00625.

## Experiment 2

### Participants

Twenty-five students (17 female, *Mage* = 21 years, ± 1.25.) at Dalhousie University participated in the current study in exchange for partial course credit. All participants had normal or corrected-to-normal vision and no history of neurological deficit as ascertained by self-report. Because all grasping movements were performed and measured using the right hand, participants were required to be comfortable performing grasping actions with their right hand. Twenty-two participants reported being right-hand dominant, one reported being left-hand dominant and two reported being ambidextrous. Each participant provided informed consent prior to participation in accordance with guidelines established by the Dalhousie University Research Ethics Board.

#### Materials

The exact same experimental setup, stimuli and procedure were used for this experiment as in experiment 1.

#### Procedure

The same procedure as in experiment 1 was applied for this experiment. The only difference was the starting position of the participants’ fingers at the start of each trial. For this experiment, participants started every trial with their right index finger and thumb spread 10.5cm apart marked by taped with tactile feedback. The position for the thumb was marked on the center of the space bar and the position for the right index finger was located on a key 10.5cm directly above the center of the space bar.

#### Data Processing

The same approach as experiment 1 was used for analysis of PGA in the GBYK and KBYG trials separately. When examining the grip aperture time course experiment 2, it was noted that early in the movement there was a decrease in grip aperture creating a local minima which we termed the local minima aperture (LMA), before the aperture increased to PGA before closing in on the object at the end of the movement (Figure 6). A similar LME was run on this LMA data as the PGA data. The Bonferroni correction was applied within outcome variable for comparisons between distractor size within condition combinations (i.e., PGA and LMA).

## RESULTS

### Experiment 1

#### GBYK

Grip aperture over the course of trial, averaged across participants within each combination of target size and distractor size is presented in Figure 4. There was a significant main effect of target size, such that smaller targets elicited reduced peak grip apertures compared to larger targets, *b* = −19.38, SE = 1.18, *t*(1974.04) = −16.49, *p* < .001. The main effect of distractor size was not significant, *b* = 0.21, SE = 1.17, *t*(1974.02) = 0.18, *p* = .856, and the effect of target location was marginal, *b* = 2.20, SE = 1.17, *t*(1974.04) = 1.88, *p* = .061. These effects were superseded by higher order interactions.

**Figure 4.**
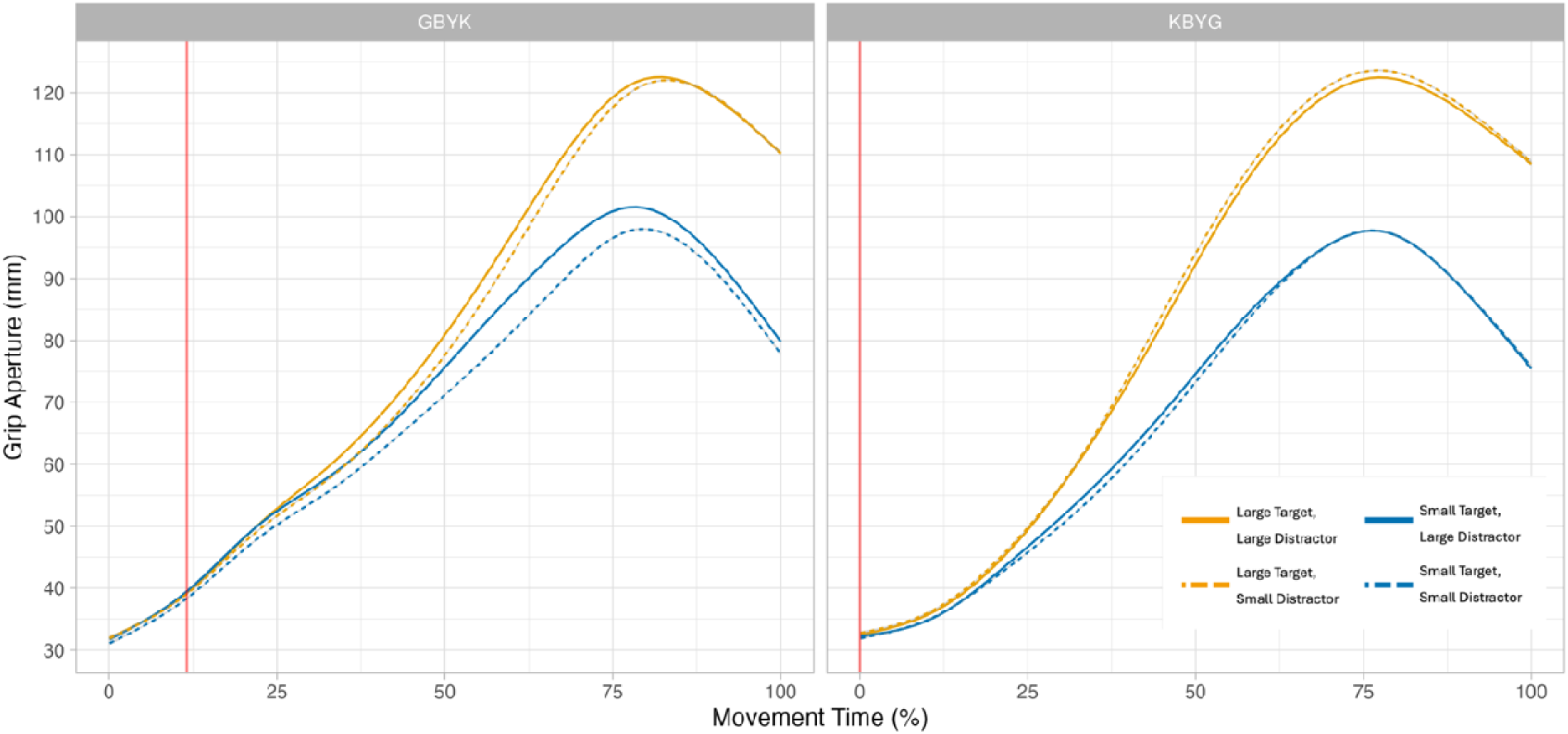
Grip aperture normalized across the trajectory in KBYG and GBYK trials for Experiment 1. The red vertical line represents target onset.

There was a significant interaction between target size and distractor size, *b* = −4.70, SE = 1.66, *t*(1974.02) = −2.83, *p* = .005, as well as a significant interaction between distractor size and target location, *b* = −3.32, SE = 1.66, *t*(1974.03) = −2.00, *p* = .045. These effects were qualified by a significant three-way interaction among target size, distractor size, and target location, *b* = 4.68, SE = 2.34, *t*(1974.02) = 2.00, *p* = .046 (Figure 5). Bonferroni corrected comparison between distractor size for each target size and location combination revealed that there was a significant difference between PGA when the distractor was large vs small when the target was small and appeared on both the left, *b* = -4.37, SE = 1.10, *t*(472.14) = 4.12, *p* <0.001, and the right side, *b* = -3.20, SE = 1.01, *t*(477.06) = 2.92, *p* =0.003. However, when the target was large, there was no impact of distractor size when the target appeared on the left, *b* = 0.164, SE = 1.21, *t*(473.10) = 0.135, *p* =0.89; and a marginal effect that did not survive the Bonferroni correct when the target was on the right, *b* = -3.10, SE = 1.20, *t*(476.12) = 2.59, *p* =0.010. Overall, these findings show that a larger distractor leads to a larger PGA compared to a small distractor consistently when the target is smaller but less consistently when the target is larger.

**Figure 5.**
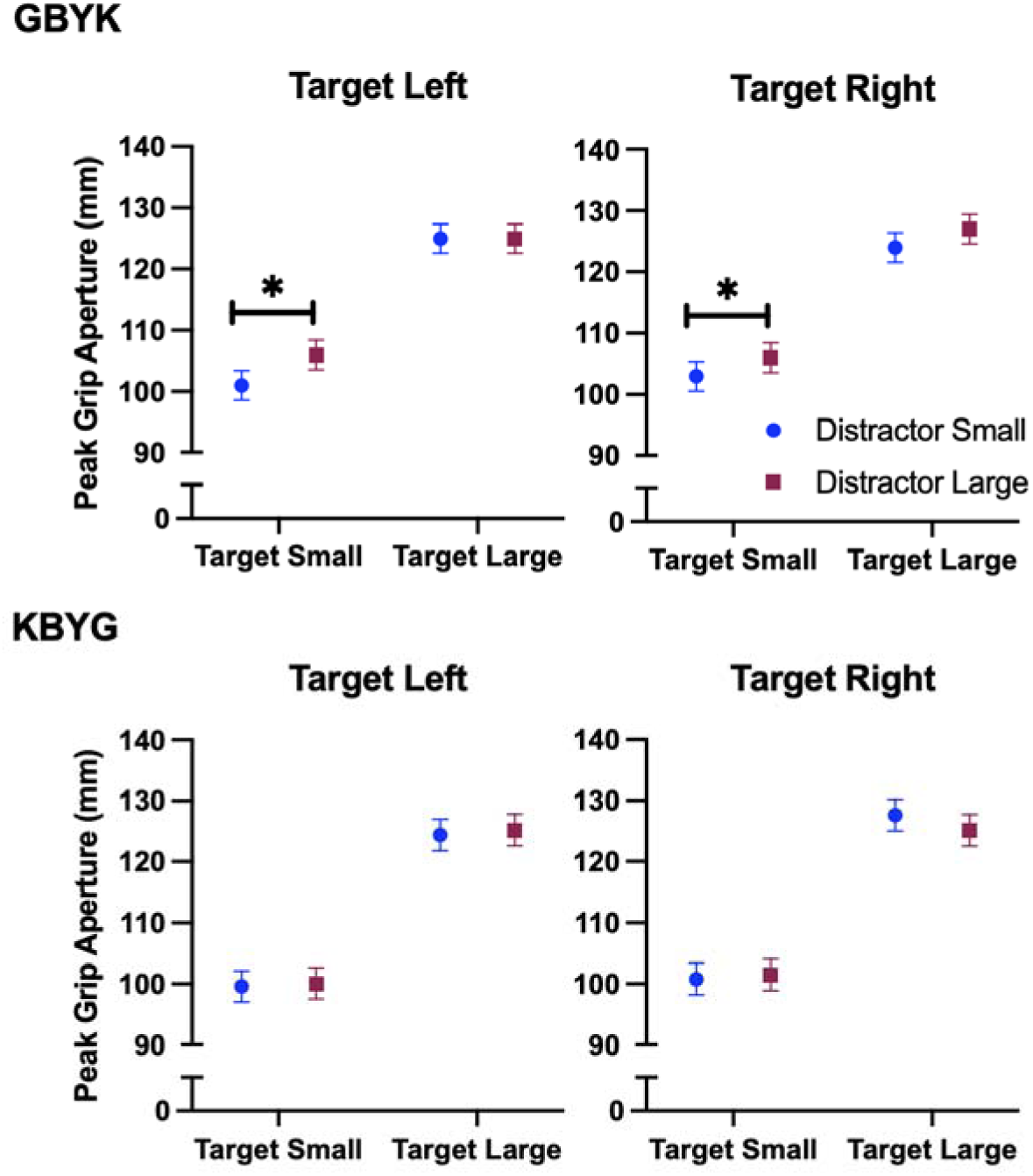
Peak grip aperture in the GBYK and KBYK trials. Error bars are standard error of the mean.

#### KBYG

There was a significant main effect of target size, where larger targets had a larger PGA compared to smaller targets, *b* = −25.11, SE = 1.27, *t*(1997.02) = −19.78, *p* < .001. The main effect of distractor size, *b* = −0.79, SE = 1.27, *t*(1997.02) = −0.62, *p* = .536, and target location, *b* = −0.14, SE = 1.27, *t*(1997.02) = −0.11, *p* = .915 were not significant (Figure 5).

None of the two-way interactions reached statistical significance, including the interaction between target size and distractor size, *b* = 0.30, SE = 1.80, *t*(1997.03) = 0.17, *p* = .865; the interaction between target size and target location, *b* = 1.53, SE = 1.80, *t*(1997.02) = 0.85, *p* = .394; the interaction between distractor size and target location, *b* = 3.37, SE = 1.80, *t*(1997.02) = 1.88, *p* = .060. The three-way interaction among target size, distractor size, and target location was not significant, *b* = −3.58, SE = 2.55, *t*(1997.03) = −1.41, *p* = .159. The planned comparisons between distractor sizes revealed no difference in PGA between distractors at any combination of the target size and location conditions [small-left: *b* = 0.56, SE = 1.23, *t*(483.10) = 0.45, *p* = .650; small-right; *b* = -0.65, SE = 1.20, *t*(1473.11) = 0.54, *p* = .59; large-left: *b* = -0.76, SE = 1.36, *t*(482.12) = 0.56, *p* = .58; large-right: *b* = 2.54, SE = 1.13, *t*(484.01) = 2.26, *p* = .02)].

### Experiment 2

#### GBYK

##### PGA

Grip aperture over the course of trial, averaged across participants within each combination of target size and distractor size is presented in Figure 6. Similar to experiment 1, smaller targets elicited reduced peak grip apertures compared to larger targets, *b* = −16.78, SE = 0.94, *t*(1893.08) = −17.80, *p* < .001. In addition, there was a significant main effect of distractor size, with smaller distractors associated with slightly reduced peak grip apertures, *b* = −1.97, SE = 0.92, *t*(1893.02) = −2.13, *p* = .034. The main effect of target location was not significant, *b* = 0.23, SE = 0.93, *t*(1893.01) = 0.25, *p* = .807. None of the two-way interactions reached statistical significance, including the interaction between target size and distractor size, *b* = −1.75, SE = 1.32, *t*(1893.03) = −1.33, *p* = .184, the interaction between target size and target location, *b* = 1.84, SE = 1.32, *t*(1893.05) = 1.40, *p* = .163, and the interaction between distractor size and target location, *b* = 1.59, SE = 1.31, *t*(1893.02) = 1.22, *p* = .223.

**Figure 6.**
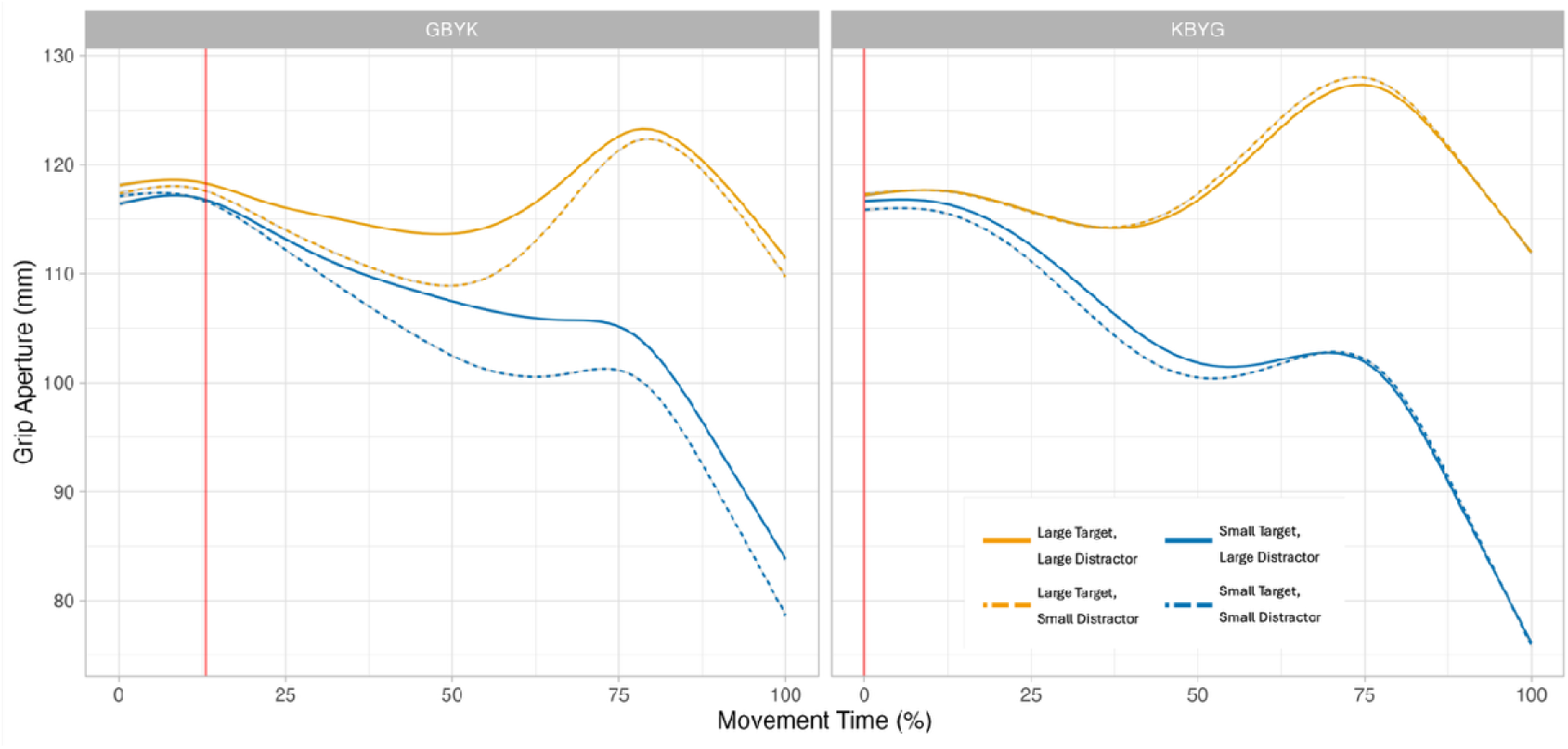
Grip aperture normalized across the trajectory in KBYG and GBYK trials for Experiment 2. The red vertical line represents target onset.

However, there was a significant three-way interaction among target size, distractor size, and target location, *b* = −4.65, SE = 1.85, *t*(1893.03) = −2.51, *p* = .012 (Figure 7) on PGA. Bonferroni-corrected comparisons between distractor sizes at each combination of target size and location revealed that there was a significant impact of distractor when the target was small on both the left, *b* = -3.74, SE = 0.80, *t*(443.28) = 4.21, *p* < .001, and right side of space, *b* = -6.70, SE = 0.94, *t*(462.05) = 7.12, *p* < .001, but not when the target was large (left side, *b* = -1.82, SE = 0.79, *t*(459,06) = 2.31, *p* = .02; right side, *b* =- 0.36, SE = 0.90, *t*(457,08) = .41, *p* = .69.

**Figure 7.**
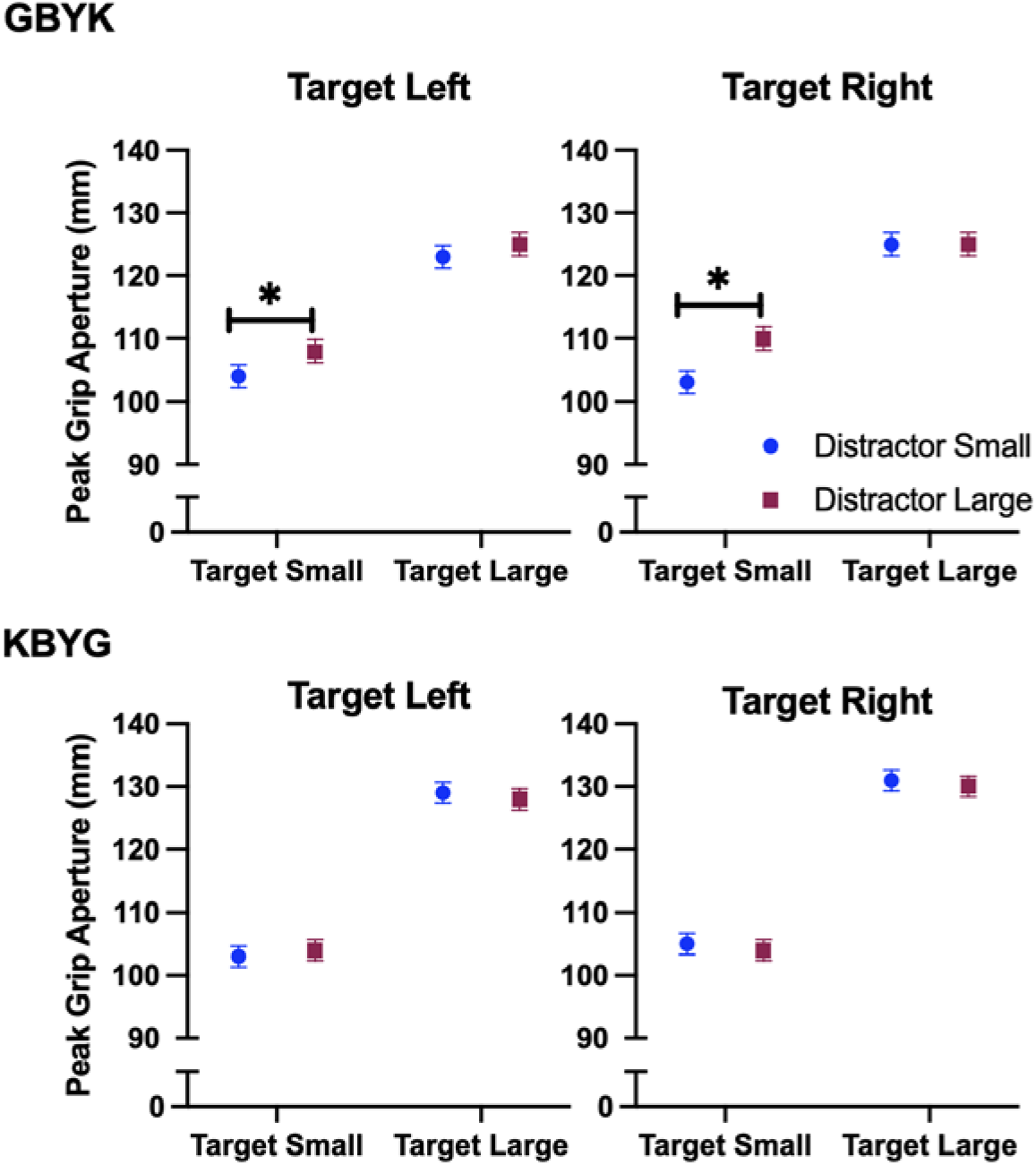
Peak grip aperture in the GBYK and KBYK trials. Error bars are standard error of the mean.

##### LMA

There was a significant main effect of target size, such that smaller targets were associated with smaller LMAs compared to larger targets, *b* = −6.92, SE = 0.78, *t*(1893.03) = −8.83, *p* < .001. There was also a significant main effect of distractor size, with smaller distractors associated with smaller LMAs, *b* = −5.83, SE = 0.77, *t*(1893.01) = −7.60, *p* < .001. The main effect of target location was not significant, *b*= 0.40, SE = 0.77, *t*(1893.00) = 0.52, *p* = .603.

None of the two-way interactions between target size and distractor size, *b* = 0.86, SE = 1.09, *t*(1893.01) = 0.79, *p* = .432, or between target size and target location, *b* = −0.78, SE = 1.10, *t*(1893.02) = −0.71, *p* = .475, were significant. However, there was a significant interaction between distractor size and target location, *b* = 2.89, SE = 1.09, *t*(1893.01) = 2.66, *p* = .008.

The three-way interaction among target size, distractor size, and target location approached significance but did not reach the conventional threshold, *b* = −2.56, SE = 1.54, *t*(1893.01) = −1.66, *p* = .096 (Figure 8); however, the planned comparisons between distractor size at each combination of target size and location were run as planned *a-priori*. These comparisons showed that that there was a significant impact of distractor when the target was small (left side, *b* = -4.99 SE = 0.79, *t*(443.13) = 6.29, *p* < .001; right side, *b* = -4.59, SE = 0.81, *t*(462.03) = 5.70, *p* < .001), and when the target was large (left side, *b* = -5.78, SE = 0.72, *t*(459.03) = 8.02, *p* < .001; right side, *b* =-2.92, SE = 0.73, *t*(457.05) = .4.02, *p* < .001).

**Figure 8.**
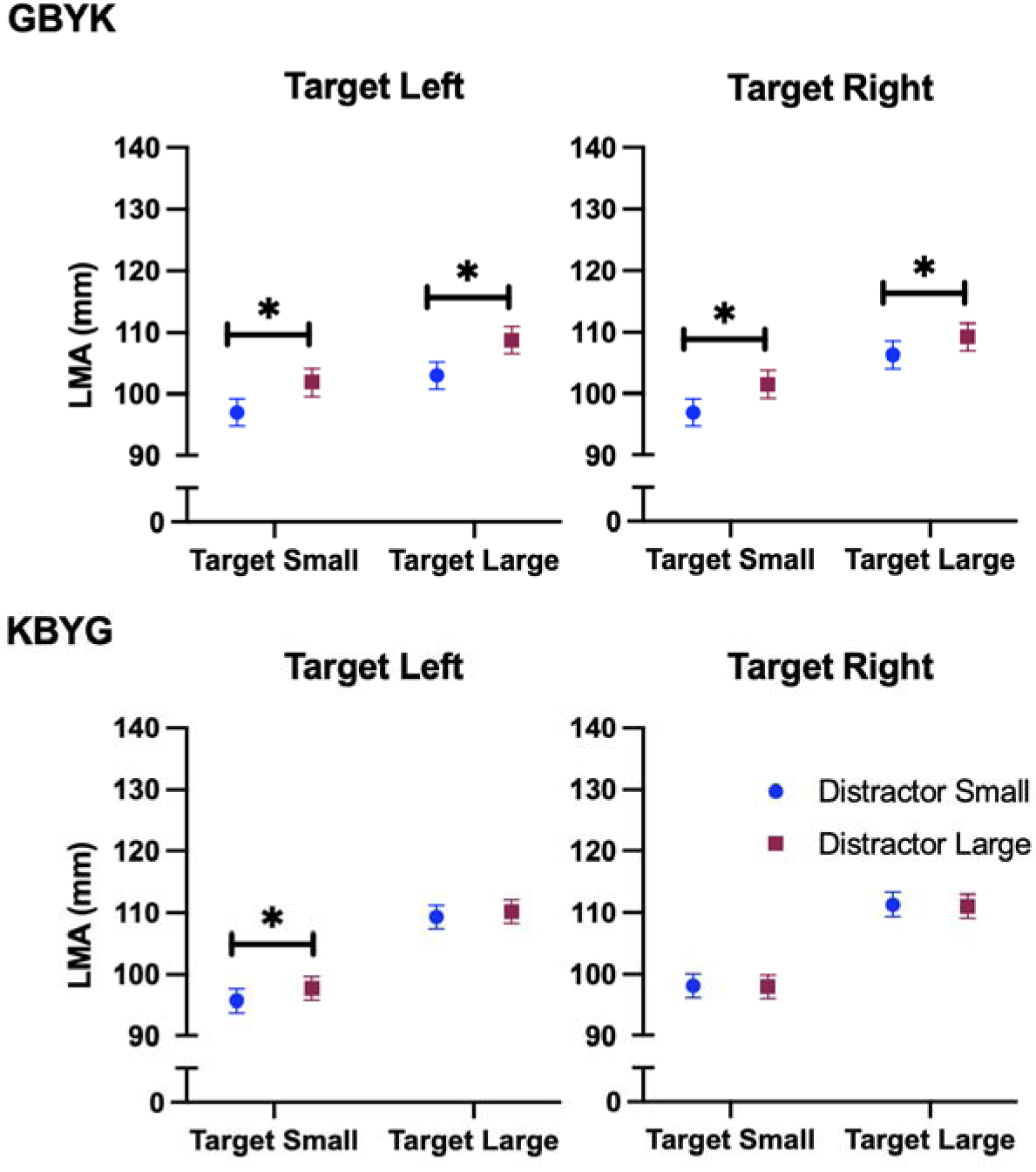
Local Minima Aperture in the GBYK and KBYK trials. Error bars are standard error of the mean.

#### KBYG

##### PGA

There was a significant main effect of target size on PGA, where small targets and a smaller PGA than lager target, *b* = −24.73, SE = 0.73, *t*(1910.00) = −33.83, *p* < .001. The main effect of distractor size was not significant, *b* = 1.04, SE = 0.73, *t*(1910.01) = 1.42, *p* = .156. There was a small but significant effect of target location, with rightward targets associated with slightly larger PGA, *b* = 1.48, SE = 0.73, *t*(1910.01) = 2.02, *p* = .044, where PGA was larger for targets on the right.

None of the two-way interactions were significant, including the interaction between target size and distractor size, *b* = −1.29, SE = 1.03, *t*(1910.01) = −1.25, *p* = .212, the interaction between target size and target location, *b* = −0.78, SE = 1.03, *t*(1910.01) = −0.76, *p* = .448, and the interaction between distractor size and target location, *b* = −0.39, SE = 1.04, *t*(1910.02) = −0.37, *p* = .709. The three-way interaction among target size, distractor size, and target location was also not significant, *b* = 1.03, SE = 1.46, *t*(1910.02) = 0.71, *p* = .480 (Figure 7). The planned comparisons between distractor sizes revealed no difference in PGA between distractors at any combination of the target size and location conditions [small-left: *b* = -0.23, SE = .75, *t*(461.00) = 0.32, *p* = .752; small-right; *b* = 0.34, SE = 0.71, *t*(463.06) = 0.49, *p* = .624; large-left: *b* = 1.05, SE = 0.69, *t*(458.98) = 1.52, *p* = .123; large-right: *b* = 0.54, SE = 0.64, *t*(455.13) = 0.84, *p* = .40)].

##### LMA

There was a significant main effect of target size on LMA, such that smaller targets had a smaller LMA compared to larger targets, *b* = −12.51, SE = 0.72, *t*(1910.00) = −17.27, *p* < .001. The main effect of distractor size was not significant, *b* = −0.93, SE = 0.73, *t*(1910.01) = −1.28, *p* = .201, and the effect of target location was also not significant, *b* = 0.84, SE = 0.73, *t*(1910.01) = 1.15, *p* = .249.

None of the two-way interactions were statistically significant, including the interaction between target size and distractor size, *b* = −1.08, SE = 1.03, *t*(1910.01) = −1.05, *p* = .293, the interaction between target size and target location, *b* = −0.60, SE = 1.02, *t*(1910.01) = −0.59, *p* = .558, and the interaction between distractor size and target location, *b* = 1.22, SE = 1.03, *t*(1910.02) = 1.19, *p* = .236. The three-way interaction among target size, distractor size, and target location was not significant, *b* = 1.01, SE = 1.45, *t*(1910.01) = 0.70, *p* = .48 (Figure 8). The planned comparisons between distractor sizes revealed there was a distractor effect for the small-left target, *b* = -2.03, SE = .69, *t*(461.00) = 2.91, *p* = .004; but not for any of the other target size/location combinations [small-right; *b* = 0.14, SE = 0.66, *t*(463.05) = 0.214, *p* = .830; large-left: *b* = -.949, SE = 0.76, *t*(459.02) = 1.25, *p* = .211; large-right: *b* = 0.16, SE = 0.63, *t*(455.03) = 0.26, *p* = .800)].

## DISCUSSION

### Experiment 1

The purpose of Experiment 1 was to determine how grasping movements are planned when two potential target objects are present and target selection occurs only after movement initiation. Specifically, we examined whether PGA, a measure of object-specific grasp planning (20–24), is influenced by the size of a nontarget object under conditions of target uncertainty. The results revealed three main findings. First, participants appropriately scaled their grip aperture according to the size of the target object, demonstrating that target-specific information was rapidly incorporated into the ongoing grasp. Second, unlike the KBYG condition, in which grip scaling was entirely unaffected by the nontarget object, PGA in the GBYK condition was influenced by the size of the nontarget. Finally, this interference was asymmetric, such that PGA was increased when reaching to grasp a small target paired with a large distractor, whereas there was minimal influence of small distractor when reaching to grasp the large target.

This pattern of result indicates that the presence of a larger nontarget object primarily influenced grasp planning when the required grip aperture was smaller, potentially reflecting competition between the motor representations of the two objects during the early stages of movement planning (50–52). In contrast, when the target object required a larger grip aperture, the smaller nontarget did not appear to influence grasp scaling. This asymmetric interference pattern is inconsistent with a motor averaging hypothesis (7–12), which predicts that distractors should produce bidirectional shifts in grip aperture toward an intermediate PGA value regardless of which object is selected as the target.

The findings are also inconsistent with alternative accounts based on precision-first processing or aperture tuning constraints. The precision-first hypothesis predicts that smaller nontarget objects should constrain grasping toward larger targets due to the increased visuomotor precision demands associated with smaller objects (58), whereas aperture tuning constraints predict that interference should depend on the initial finger configuration (59). Instead, the opposite pattern was observed: a large distractor influenced grasping toward a small target, whereas a small distractor did not influence grasping toward a large target.

The results are most consistent with a safety-based optimization hypothesis in which, under conditions of uncertainty, participants appear to prioritize information from the larger object when present, increasing PGA beyond that required for the small target when paired with a larger distractor while avoiding interference when the target already requires a larger aperture. Thus, when multiple potential targets are present, grasp planning appears to incorporate nontarget information in a manner that prioritizes successful object acquisition (i.e., motor optimization mechanism, 13-18) rather than averaging competing representations or other mechanisms. Specifically, opening the hand slightly wider than necessary when reaching for a small object is unlikely to compromise successful acquisition, whereas selecting an aperture that is too small for a larger object could increase the risk of an unsuccessful grasp (e.g., tipping the object during contact). Therefore, under conditions of target uncertainty, the motor system may adopt a conservative grip scaling strategy that minimizes the likelihood of grasp failure by incorporating potentially relevant information from larger competing objects.

Experiment 2 was originally designed to dissociate between two potential explanations if a nontarget interference effect from the small object was found: whether such interference reflected increased online processing of competing object representations or constraints arising from overlapping grip aperture tuning. By altering the initial finger positioning, Experiment 2 was intended to determine whether nontarget interference from the smaller object depends on the starting state of the fingers. However, because Experiment 1 did not reveal a nontarget interference via the small object and instead demonstrated an asymmetric effect in which a large object nontarget increased PGA when grasping a small target, Experiment 2 was repurposed to replicate this pattern and rule out initial finger position as a potential explanation. If the observed interference reflects a safety-based optimization strategy, changing the starting position of the fingers should not alter the pattern of results. Specifically, initiating movements with the fingers apart should still produce larger PGA when grasping a small target paired with a large distractor, with no influence of a small distractor when grasping a large target. Conversely, if the asymmetric interference observed in Experiment 1 was driven by constraints associated with the initial aperture of the fingers (although not in the direction originally predicted), altering the starting configuration should change the pattern of interference.

### Experiment 2

The purpose of Experiment 2 was to determine whether the asymmetric nontarget interference observed in Experiment 1 depended on the initial start position of the fingers. Despite participants initiating movements with their fingers apart, the pattern of PGA results replicated those of Experiment 1. Participants appropriately scaled their grip aperture to the target object, and PGA was again selectively influenced by the presence of a larger nontarget object when grasping the small target, with no effect of a small nontarget when grasping the large target under the GBYK condition. This replication indicates that the asymmetric interference observed in Experiment 1 cannot be attributed to constraints associated with the starting aperture of the fingers. Instead, the consistency of the effect across experiments further supports the interpretation that under conditions of target uncertainty, grasp planning incorporates information from larger competing objects in a manner consistent with a safety-based optimization approach to maximize object acquisition.

However, interestingly, analysis of the grip aperture time course revealed an additional feature that was not apparent in Experiment 1. Early in the movement, grip aperture initially decreased, producing a local minima aperture (LMA) before subsequently increasing to PGA. Unlike PGA, LMA exhibited evidence of nontarget interference consistent with an intermediate aperture response predicted by the motor averaging hypothesis (9–12). Specifically, in the GBYK condition, the aperture at this early stage of the movement was biased by the size of the competing object, suggesting that representations of both potential targets may have contributed to the initial stages of movement planning. These findings suggest that information about both potential target objects is incorporated during the earliest stages of grasp planning, producing an initial grip aperture that reflects the influence of competing object representations. This pattern is consistent with a motor averaging account, although it could also reflect a strategic optimization process in which participants adopt an intermediate aperture to maximize the likelihood of successfully grasping either potential target under conditions of uncertainty. Distinguishing between these explanations will require further investigation. However, as the movement unfolds, this early averaging/intermediate process is no longer evident, and the final grip aperture instead reflects the safety-based optimization pattern observed at PGA.

Importantly, this dissociation emerged only when movements were initiated with the fingers apart in Experiment 2. In contrast, the grip aperture trajectory observed in Experiment 1 followed the typical pattern reported across previous grasping studies (2–6), characterized by a smooth increase in aperture toward PGA followed by closure of the fingers around the object – no distinct pre-kinematic pattern prior to PGA. Thus, the wide starting aperture used in Experiment 2 may have provided a distinct point from which to examine the early portion of the grasping movement, allowing us to consider whether early movement reflected competition between potential object representations, whereas later movement reflected target-specific optimization. Moreover, there was some evidence distractor interference in LMA even under the KBYG condition, despite previous evidence that grip aperture during visually guided grasping is generally resistant to distractor effects (Exp 1, 53-54). This finding suggests that initiating movements with the fingers apart may have altered the natural dynamics of grasp planning. Some caution should be used in interpreting this result because it only emerged in one of our planned comparisons and not in the main model. Nevertheless, prior work has shown that unnatural or consciously monitored movements are more susceptible to perceptual interference, potentially due to increased interactions between dorsal and ventral stream processing (60). The potential implications of these findings, as well as other possible factors that may contribute to this pattern, such as object weight, are considered further in the General Discussion.

## GENERAL DISCUSSION

The primary finding of the present study is that under conditions of target uncertainty, grasp planning appears to prioritize successful object acquisition rather than integrating competing object representations to produce an intermediate grip aperture. Across both experiments, PGA was appropriately scaled according to the target object, demonstrating that object-specific information was successfully incorporated into the grasping movement despite uncertainty regarding target identity. However, unlike the KBYG condition, where grasp scaling was entirely determined by the target object, PGA in the GBYK condition was influenced by the presence of a larger nontarget object. This influence was asymmetric, such that a larger nontarget increased PGA when grasping a smaller target, whereas a smaller nontarget had no effect when grasping a larger target. This pattern is inconsistent with a motor averaging mechanism (9–12), which would predict that competing object representations should bias grip aperture toward an intermediate value regardless of which object becomes the target. Instead, the results suggest that when multiple potential targets are present, the visuomotor system incorporates competing object information in a manner that prioritizes minimizing the likelihood of grasp failure (i.e., movement optimization: 13-18).

This safety-based optimization account is consistent with broader theories of motor planning suggesting that actions are selected based on their expected consequences rather than simply reflecting competition between simultaneously activated motor representations (13). When uncertainty exists regarding the identity of the target object, selecting a slightly larger grip aperture represents an advantageous strategy because it maintains successful acquisition across possible outcomes. Specifically, opening the hand wider than required for a small object is unlikely to prevent successful grasping, whereas selecting an aperture that is too small for a larger object increases the risk of unsuccessful contact or object displacement. Thus, under conditions of uncertainty, the grasping system may bias motor planning toward solutions that maximizes acquisition rather than precision.

The results also provide evidence against precision-first and overlapping aperture tuning accounts, both of which predict that interference should be driven by the smaller object. Specifically, these accounts would predict reduced PGA when grasping a larger target paired with a small nontarget, either because smaller objects impose greater visuomotor precision demands (58) or because aperture scaling is constrained by the initial finger configuration (59). However, across both experiments, we observed the opposite pattern: larger nontarget objects increased PGA when grasping smaller targets, whereas smaller nontargets did not influence PGA when grasping larger targets. Given that PGA reflects how the hand is scaled to match object size during grasp formation (2–6), these findings suggest that target uncertainty does not simply constrain grasp aperture based on the most precise or restrictive object but instead reflects a bias toward larger object representations when multiple potential targets are present.

The present findings also suggest that the influence of competing object representations can emerge under certain movement contexts. In Experiment 2, when participants initiated the movement with their fingers separated, analysis of the grip aperture trajectory revealed evidence that the earliest stages of the movement were influenced by both potential targets. Specifically, the local minima aperture (LMA) showed a pattern consistent with integrating competing object representations to produce an intermediate grip aperture, such that the initial aperture reflected an integration of the competing object size representations. This finding suggests that although the final grasp configuration is optimized according to task demands, competing motor representations may still contribute during the initial stages of movement formation.

Importantly, however, this early averaging-like influence should be interpreted cautiously, as the movement context in Experiment 2 differed from typical visually guided grasping tasks. In addition to the presence of target uncertainty, participants initiated movements from a starting posture in which the thumb and fingers were widely separated. Although this configuration may more closely resemble natural situations in which the hand is open and ready to interact with objects, it differs from the closed or partially closed starting postures commonly used in experimental grasping studies. Maintaining a wide finger separation prior to movement may also introduce additional biomechanical demands, requiring participants to actively maintain the hand in an extended configuration against its natural resting posture. These factors may have altered the dynamics of grasp formation and increased susceptibility to interference from competing object representations. Consistent with this interpretation, nontarget interference was also observed in the KBYG condition of Experiment 2 (e.g., a small target presented to the left was grasped with a larger PGA when a large distractor was presented to the right), despite participants having full knowledge of the target object prior to movement initiation. Since previous research has shown that grip aperture during typical visually guided grasping is highly resistant to distractor interference (Experiment 1; 53-54), the presence of distractor effects even when the target was known suggests that the observed influence may not solely reflect target uncertainty but may also reflect the unique biomechanical demands associated with the starting configuration.

There is a possibility that initiating movements from a wide starting aperture increased competition between simultaneously activated grasp representations, making the movement more vulnerable to interference from alternative motor commands. Alternatively, the altered movement context may have increased reliance on perceptual processing mechanisms associated with conscious monitoring of the action. Previous work has suggested that movements performed under less automatic or explicitly controlled conditions can increase interactions between dorsal and ventral visual stream processes, potentially allowing perceptual object representations to exert greater influence over visually guided actions (60). Consistent with this possibility, LeBlanc and Westwood (61) demonstrated that a preceding perceptual task influenced grip aperture scaling during a grasping action, suggesting that perceptual processing can contribute to the online shaping of grasp kinematics. Therefore, the early averaging-like LMA effects observed in Experiment 2 may reflect either increased competition between motor representations or greater susceptibility to perceptual interference under conditions where the typical automaticity of grasp control is reduced by the unique biomechanical demands of the starting configuration.

Another important consideration is that object size was confounded with other object properties, including mass and visual salience. The larger object was 95 g heavier than the smaller object, meaning that the influence of a larger nontarget object on PGA may reflect not only adjustments based on spatial requirements for hand shaping but also expectations regarding the force demands associated with grasping a heavier object. Indeed, recent neuroimaging evidence suggests that while object size and grasp axis are primarily represented during grasp planning within dorsal-stream visuomotor regions, object mass is incorporated more strongly during movement execution through interactions between ventral-stream and premotor regions involved in grip force regulation (29). Additionally, larger objects may receive greater attentional weighting due to their increased visual salience (62–64). Therefore, the increased PGA observed when a larger nontarget object was present may reflect a combination of spatial, attentional, and force-related influences rather than object size scaling alone.

Future studies should isolate the contribution of object size by manipulating it independently from other object properties. For example, rectangular objects with identical mass and dimensions but different orientations could vary the required grasp aperture while controlling for weight and visual salience. This approach would allow researchers to determine whether the optimization observed under target uncertainty primarily reflects sensitivity to grasp size and hand-shaping requirements or the integration of additional object properties, such as weight, during action planning.

In conclusion, the present study provides novel evidence that grasp planning under target uncertainty involves the integration of competing object information in a manner that prioritizes successful object acquisition. Across two experiments, participants were able to appropriately scale grip aperture according to the target object despite initiating movements before target identity was known. However, when multiple potential targets were present, PGA was influenced by the presence of a larger nontarget object, such that a larger distractor increased aperture when grasping a smaller target, whereas a smaller distractor had no influence when grasping a larger target. This asymmetric pattern of interference is inconsistent with a motor averaging mechanism, which would predict that competing object representations bias grip aperture toward an intermediate value regardless of which object becomes the target. Instead, these findings suggest that under conditions of uncertainty, grasp planning incorporates information from competing objects in a manner that prioritizes motor solutions that maximize the likelihood of successful object acquisition. Although early movement dynamics under altered starting conditions revealed evidence that competing representations can influence the initial stages of grasp formation, the final grasp configuration appears to reflect a task-dependent optimization process. This study extends previous work on target uncertainty action selection beyond reaching, eye, and stepping movements and highlights that grasp planning under uncertainty is a dynamic process in which competing object representations are refined into a motor plan optimized for successful object acquisition.

## DATA AVAILABILITY

Data are available on https://doi.org/10.5683/SP4/5OITGH

## AUTHOR CONTRIBUTIONS

KAL and HFN conceived and designed research; ATDL, JAB, and MG performed experiments; ATDL, BBTF, and HFN analyzed data; KAL and HFN interpreted results of experiments; ATDL, JAB, and HFN prepared figures. KAL, ATDL, JAB, and HFN drafted the manuscript; All authors reviewed and edited a final draft of the manuscript. KAL and HFN approved final version of manuscript.

